# Inhibition is a prevalent mode of activity in the neocortex around awake hippocampal ripples

**DOI:** 10.1101/2022.03.02.482726

**Authors:** Javad Karimi Abadchi, Zahra Rezaei, Thomas Knöpfel, Bruce L. McNaughton, Majid H. Mohajerani

## Abstract

Coordinated peri-ripple activity in the hippocampal-neocortical network is essential for mnemonic information processing in the brain. Hippocampal ripples likely serve different functions in sleep and awake states. Thus, the corresponding neocortical activity patterns may differ in important ways. We addressed this possibility by conducting voltage and glutamate wide-field imaging of the neocortex with concurrent hippocampal electrophysiology in awake mice. Contrary to our previously published sleep results, deactivation and activation were dominant in post-ripple neocortical voltage and glutamate activity, respectively, especially in the agranular retrosplenial cortex (aRSC). Additionally, the spiking activity of aRSC neurons, estimated by two-photon calcium imaging, revealed the existence of two subpopulations of excitatory neurons with opposite peri-ripple modulation patterns: one increases and the other decreases firing rate. These differences in peri-ripple spatiotemporal patterns of neocortical activity in sleep versus awake states might underlie the reported differences in the function of sleep versus awake ripples.

## Introduction

Hippocampal-neocortical interactions around hippocampal ripples play an important role in memory processes^1–4^. The functional role of such interactions are believed to be brain state-dependent such that they are involved in memory consolidation during NREM sleep, while they are implicated in memory-guided behavior such as planning and memory retrieval in waking state^5–7^. This state-dependent functional dichotomy poses a question: how does spatiotemporal dynamics of hippocampal-neocortical network interactions differ in the two states?

There are pronounced differences in the neocortical activity patterns between NREM sleep and quite wakefulness. The most prominent difference is the near absence of so-called slow-oscillations (SO) during wakefulness. SO, a quasi-synchronous <= 1 Hz rhythmic fluctuation observed in LFP and EEG recordings throughout the neocortex during NREM sleep, is partly correlated with hippocampal ripples, and recent memory reactivation in cortex is strongly locked to ripples^8–11^. However, given the near absence of SO in wakefulness^12^, it is not straightforward to extrapolate from sleep to wakefulness, although a few studies have shown that the proportion of neurons whose spiking activity is suppressed around hippocampal ripples is significantly higher in wakefulness compared with sleep.

In the present study, we extended the previous results by imaging the activity of a large portion of the dorsal neocortical mantle in awake mice, with concurrent local-field potential (LFP) and multi-unit activity (MUA) recording from the pyramidal layer of the dorsal CA1. Wide-field glutamate and voltage recording were used to capture the excitatory synaptic input and the membrane potential fluctuations across neocortical regions and to correlate them with the occurrence of hippocampal ripples. A sharp contrast in the peri-ripple neocortical activity between the awake and sleep states was observed. To further elaborate on this contrast, we used the two-photon calcium imaging to focus on the agranular retrosplenial cortex (aRSC) whose glutamate and voltage activity patterns were different from the rest of the imaged regions. Our results suggest that inhibition is more pronounced in peri-ripple neocortical activity in awake than sleep states.

## Results

To study peri-ripple activity across neocortical regions in the awake state, we utilized three imaging modalities to shed light on different aspects of the problem at hand. First, to capture the internal dynamics of neocortical regions, wide-field voltage imaging with voltage indicator (butterfly1.2; VSFP mice) expressed in the excitatory neurons of the neocortical layers II/III was used (Fig. 1aii). Second, to capture the excitatory input to the neocortical regions, wide-field imaging with intensity-based glutamate-sensing fluorescent reporter (iGluSnFR; iGlu-Ras mice) indicator expressed in the excitatory neurons of the neocortical layers II/III was used (Fig. 1ai). Last, to estimate the spiking output of neocortical neurons, two-photon calcium imaging of the aRSC superficial layers in Thy1-GCamp mice was conducted (Fig. 1aiii). In addition, to compare the peri-ripple glutamatergic transmission in superficial versus deep neocortical layers, wide-field imaging of iGluSnFR activity in all neocortical layers (iGlu-EMX mice) was conducted. In all the imaging modality experiments, concurrent local field potential (LFP) and multi-unit activity (MUA) recordings from pyramidal layer of the CA1 subfield of the dorsal hippocampus was performed, and the hippocampal LFP was used to detect ripples (Fig. 1b). Moreover, in 3 out 4 sets of the experiments, electromyography form the neck muscles was conducted to monitor the animals’ movements. During the recordings, the mice were placed on a stationary platform in all the experiments to increase the probability of occurrence of motionless periods during which ripples would emerge. In general, the animals did not move right before the ripples; however, they sometimes tend to move after ripples occurred (Supplementary Fig. 1). Hence, to remove the potential role of body movement on the brain optical signals, ripples with EMG tone around them (±500 ms) were excluded (0.2847 ± 0.1576; mean ± std; n = 19 animals). Then, to extract the overall neocortical activity around ripples, the activity, captured by the three modalities, was aligned with respect to the timestamp of the ripple centers (largest trough of ripples) and averaged (Figure 1c). In addition, to evaluate the coordination between hippocampal and neocortical activity, the ensemble-wise correlation coefficient of the hippocampal MUA and the activity of neocortical regions were calculated (see Methods).

**Fig. 1.**
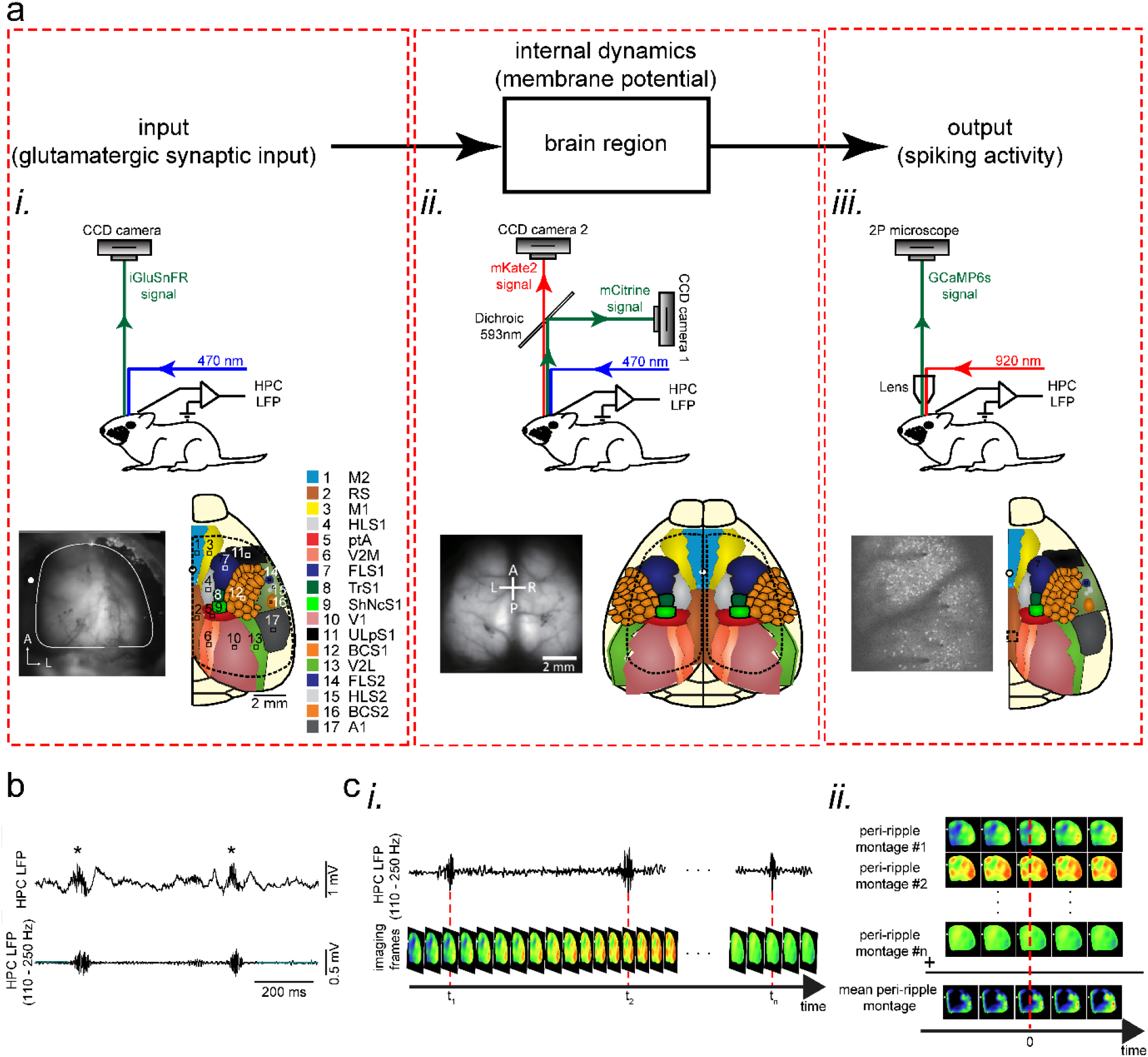
Experimental protocol for investigating peri-ripple neocortical activity during awake state. **a** Top: each region could be modeled as an input-output block with internal dynamics. Bottom (i-iii): experimental setups, exemplar imaging windows, and schematic of the regions included in the windows for unilateral wide-field glutamate imaging (i), bilateral wide-field voltage imaging (ii), and two-photon calcium imaging (iii) which were conducted for monitoring input, internal dynamics, and output, respectively. **b** Top: an exemplar local field potential (LFP) trace recorded from pyramidal layer of CA1 subfield of the dorsal hippocampus. Asterisks denote detected ripples. Bottom: ripple-band (110-250 Hz) filtered version of the top trace. **c** Schematic of peri-ripple (ripple-triggered) averaging analysis. (i) Schematic of concurrently recorded LFP and imaging signals. Red dashed lines indicate timestamp of center of detected ripples. (ii) The imaging frames around the timestamp of the detected ripples are aligned with respect to the ripple centers and averaged.

**Supplementary Fig. 1.**
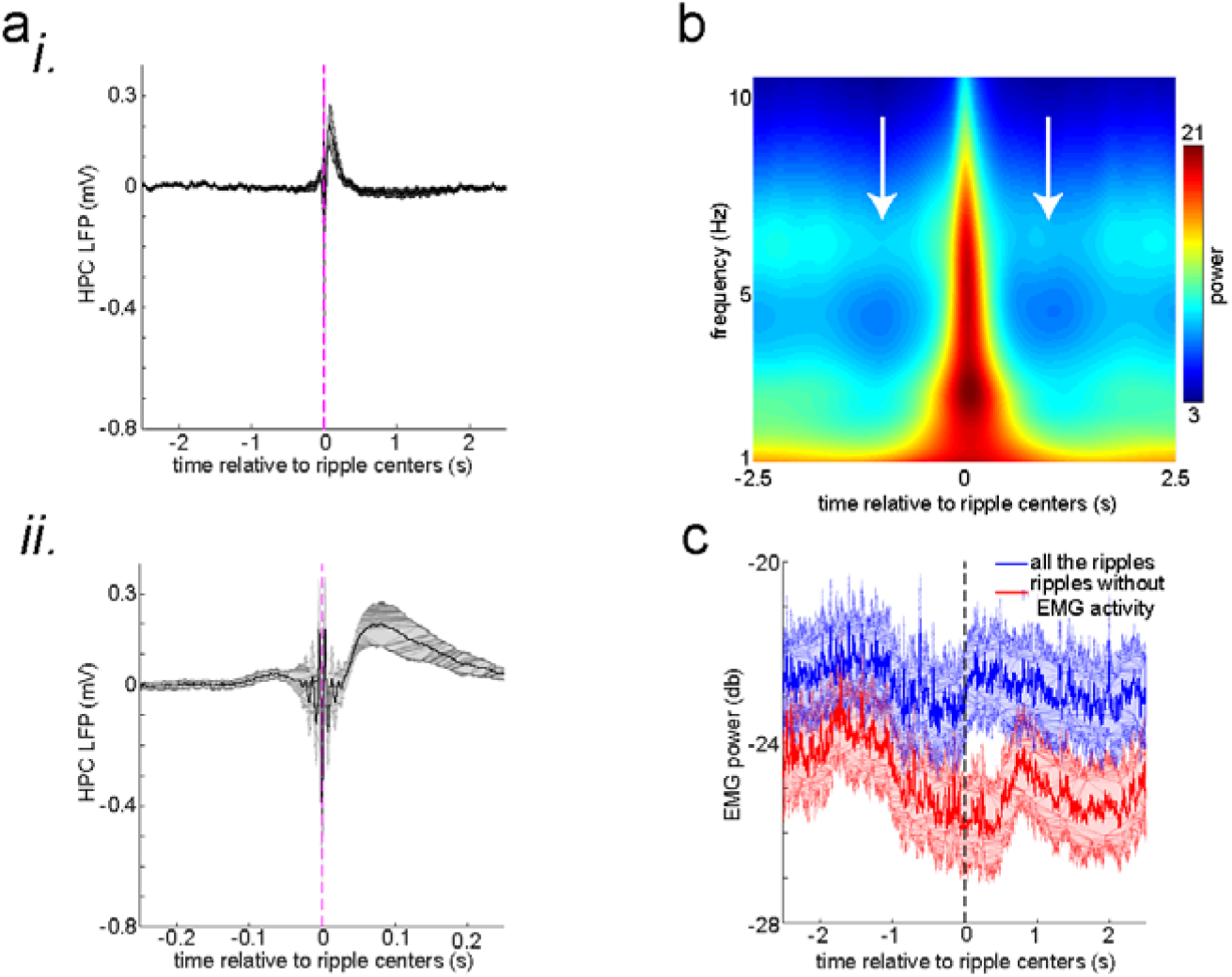
**related to Fig. 1 a (i)** The **m**ean peri-ripple hippocampal local field potential (HPC LFP) first averaged in individual animals and then averaged across 25 animals. **(ii)** The zoomed-in version of (i). **b** The mean spectrogram of peri-ripple HPC LFP first averaged in individual animals and then averaged across 25 animals. Note the high power of ∼3 Hz close to 0 time corresponding to the post-ripple large deflection apparent in (a). Moreover, note the reduction of theta power (∼6 Hz; white arrows) before and after the ripple centers time (0 time). **c** The **m**ean peri-ripple EMG signal first averaged in individual animals and then averaged across 19 animals. The blue trace is associated with all the detected ripples, and the red trace is associated with all the ripples around which (±500 ms) no EMG activity was detected. Note that both traces show a significant reduction right before the ripple centers (0 time). Note also that the blue trace shows an elevation right after the ripple centers indicating that the animals start moving right after a proportion of ripples (0.2847 ± 0.1576; mean ± std; n = 19 animals). The shadings represent the standard error of the mean.

### Population membrane voltage significantly dropped during awake ripples in the neocortical superficial layers

The ripple event-triggered averaged neocortical membrane voltage showed a fast hyperpolarization right after the ripple centers (Fig. 2ai, 2bi-iii). These peri-ripple voltage signals in the awake state were in sharp contrast with what had been reported in sleep where a membrane depolarization dominates^13^ (Supplementary Fig. 2). There was significant regional variation in the hyperpolarization pattern, with aRSC showing the strongest reduction of amplitude (Fig. 2b). This phenomenon was consistent in all the six animals used for this set of experiments (Supplementary Fig. 3a). The rate of reduction of voltage was also fastest in aRSC in the majority of animals (Supplementary Fig. 3b). Moreover, we observed a pre-ripple elevation of voltage which was also strongest in aRSC in at least half of the animals (Fig. 2biii; Supplementary Fig. 3c). Lastly, the ensemble-wise correlation coefficients averaged across VSFP animals revealed a period (∼ 0 – 100 ms) of enhanced coordination between hippocampal MUA and aRSC voltage activity which was absent for somatosensory regions (Fig. 2biv-vi).

**Fig. 2.**
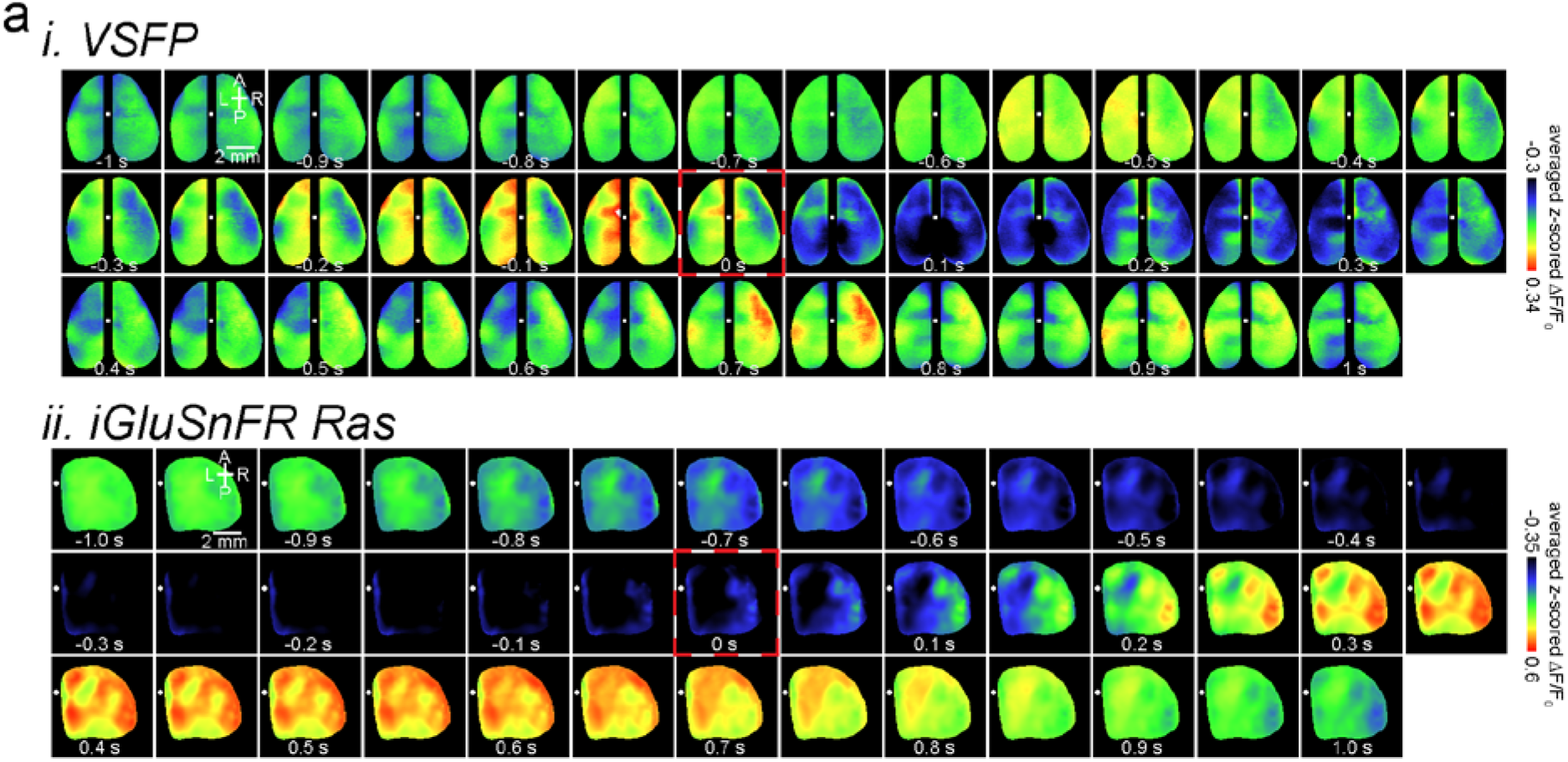
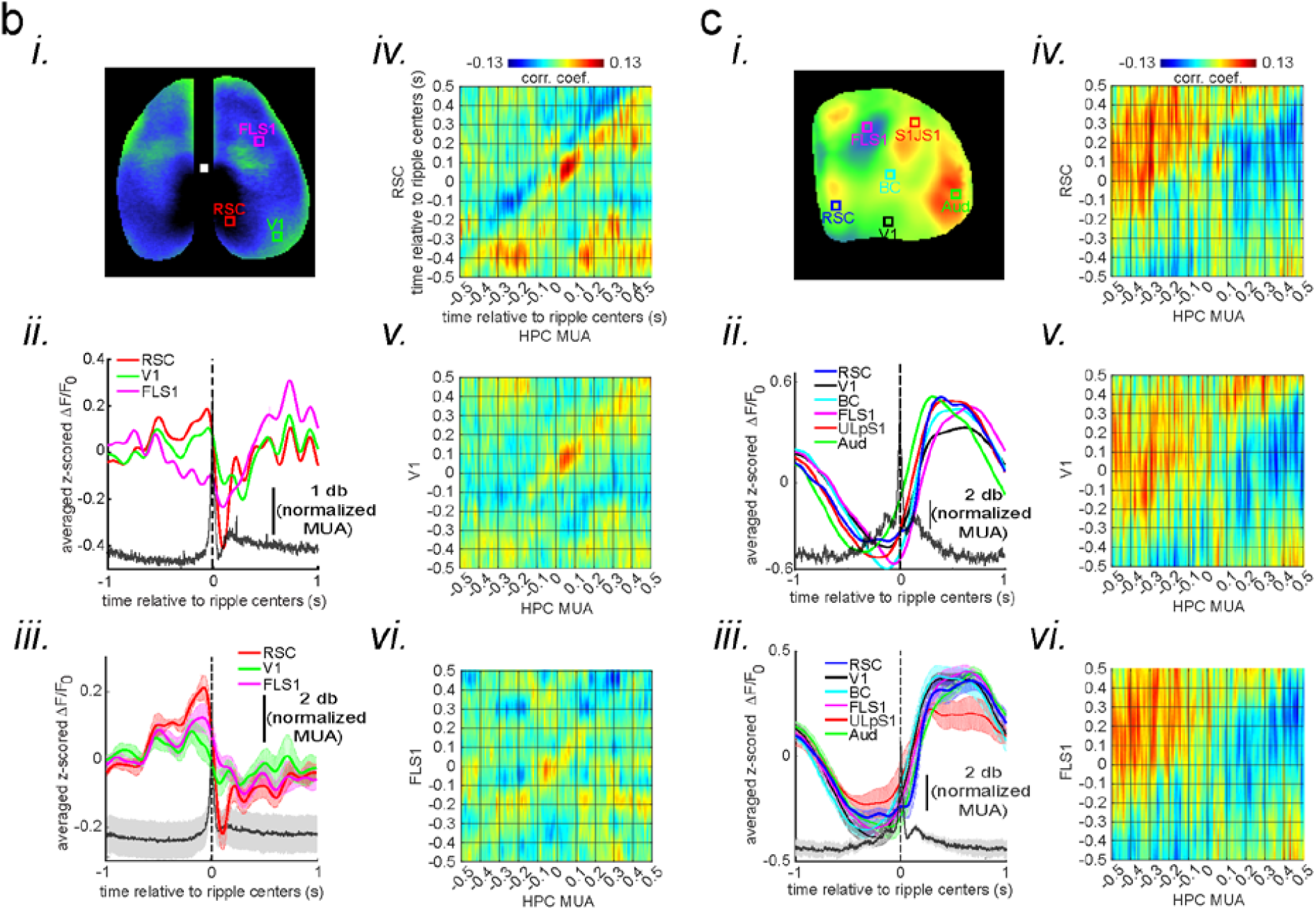
Deactivation and activation dominate the neocortical voltage and glutamate activity, respectively, during awake ripples. **a (i-ii)** Montage of average voltage (i) and glutamate (ii) activity 1 second before and after ripple centers in two representative animals. Zero time (red dashed square) represents the timestamp of center of ripples. Note the reduction of voltage signal across neocortical regions during ripples and the elevation of voltage activity before ripples. The deactivation is strongest in the agranular retrosplenial cortex (aRSC), the dark area in the posterior-medial part the imaging window which is noticeable in the frame associated with the time 100 ms in (i). Glutamate activity, on the other hand, showed a strong activation during ripples in all the regions. **b (i-ii)** A representative frame chosen from hyperpolarization period in (a-i) along with peri-ripple mean voltage time-series of three region of interests chosen from agranular retrosplenial cortex (aRSC), primary visual cortex (V1), and primary forelimb somatosensory cortex (FLS1). The data represented in time-series format is the same data shown in (a-i). The black trace represents the mean hippocampal multi-unit activity (HPC MUA). **(iii)** Peri-ripple mean voltage and MUA time-series averaged across six VSFP mice. The shading represents standard error of the mean (SEM). aRSC shows strongest and fastest deactivation compared with other regions. **(iv-vi)** Ensemble-wise correlation coefficient function of the peri-ripple voltage activity of the neocortical regions and hippocampal multi-unit activity (MUA). Rows and columns of the matrices represent time (in seconds) relative to ripple centers. **(c)** The same as (b) but for iGlu-Ras animals (n = 4) with extra ROIs from primary lip somatosensory cortex (ULpS1), primary barrel cortex (BC), and primary auditory cortex (Aud). The glutamate signal from aRSC shows fastest and latest onset of elevation. Note the presence and absence of enhanced correlation between aRSC and HPC MUA in the time interval (0,100 ms) in the voltage and glutamate activity, respectively.

**Supplementary Fig. 2.**
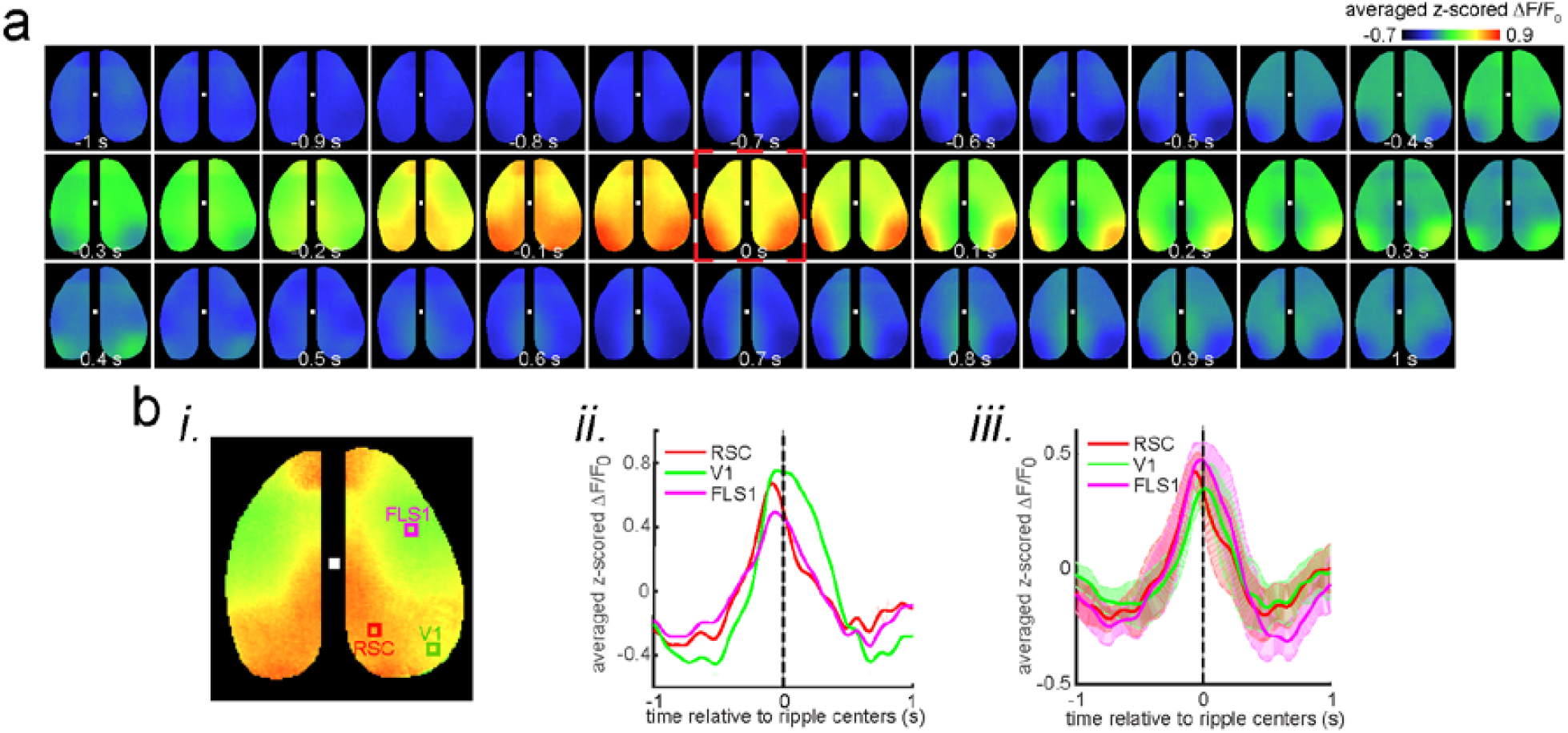
**related to Fig. 2 a** Montage of average voltage activity 1 second before and after ripple centers under urethane anesthesia (as a model of sleep) in the same representative animal as in Fig. 2a-i. Zero time (red dashed square) represents the timestamp of center of ripples. Note the elevation of voltage signal (depolarization) across neocortical regions during ripples which is in sharp contrast with the result in Fig. 2a-i. **b (i-ii)** A representative frame chosen from depolarization period in (a) along with peri-ripple mean voltage time-series of three region of interests chosen from agranular retrosplenial cortex (aRSC), primary visual cortex (V1), and primary forelimb somatosensory cortex (FLS1). The data represented in time-series format is the same data shown in (a). **(iii)** peri-ripple mean voltage time-series under urethane anesthesia averaged across five VSFP mice. The shading represents standard error of the mean (SEM).

**Supplementary Fig. 3.**
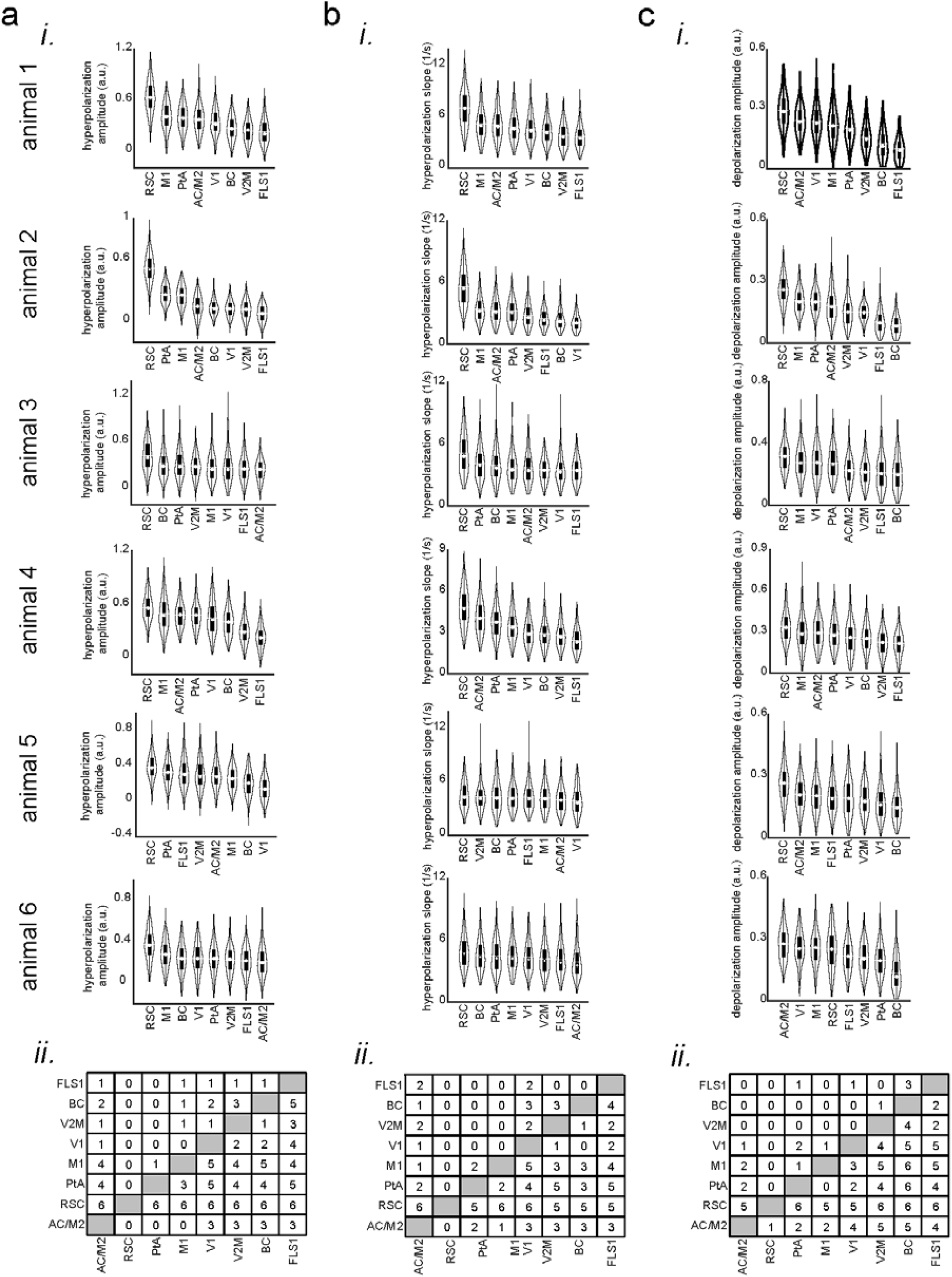
**related to Fig. 2 a (i)** Bootstrap distribution of voltage reduction amplitude (with respect to the pre-ripple baseline) across imaged neocortical regions for individual VSFP animals. The regions are sorted according to their distribution mean in descending order (repeated-measure ANOVA with df = 7 and n = 231; from top to bottom: F = 440.597, p-value = 1.0596×10^−211^; F = 858.17, p = 0; F = 62.95, p = 3.0723×10^−46^; F = 202.86, p = 1.578×10^−86^; F = 163.62, p = 1.797×10^−82^; F = 115.303, p = 2.703×10^−97^). **(ii)** The result of posthoc multiple comparisons following repeated-measure ANOVA, pooled across animals (6 animals). The numbers at each entry of the matrix represent the number of mice for which the bootstrap distribution associated with the corresponding row region has statistically significant larger mean than column region. Significance alpha level of 0.05 was used as the threshold for statistical significance. **b-c** The same as (a) but for voltage reduction slope (mean over full-width at half-maximum of derivative of the voltage signal; repeated-measure ANOVA with df = 7 and n = 231; from top to bottom: F = 195.096, df = 7, p-value = 1.1431×10^−128^; F = 291.61, p = 1.32×10^−161^; F = 63.9, p = 4.56×10^−55^; F = 240.13, p = 1.36×10^−146^; F = 7.35, p = 4.06×10^−6^; F = 20.24, p = 1.43×10^−20^) and pre-ripple voltage elevation amplitude (mean over full-width at half-maximum; repeated-measure ANOVA with df = 7 and n = 231; from top to bottom: F = 288.5488, df = 7, p-value = 1.8389×10^−142^; F = 316.86, p = 1.79×10^−213^; F = 92.71, p = 4.53×10^−74^; F = 62.03, p = 4.88×10^−37^; F = 58.77, p = 2.3×10^−50^; F = 134.77, p = 2.37×10^−123^). Note that aRSC, compared with all other imaged regions, shows largest voltage reduction amplitude, fastest rate of change of voltage reduction, and largest pre-ripple voltage elevation in at least 5 out of 6 animals.

### Glutamate concentration increases after awake ripples in the neocortical superficial layers

Next, we performed peri-ripple averaging of glutamate indicator (iGluSnFR) signals of the neocortical superficial layers. In all imaged neocortical regions, the glutamate signal was reduced before the ripple peak (Fig. 2aii, ci-iii). This reduction is probably associated with a brain state (i.e., quiet wakefulness) which is conducive for emergence of the ripples (Supplementary Fig. 1). On the other hand, after the ripples occurred, the glutamate signal increased (Fig. 2aii, ci-iii). The amplitude of the signal varied between regions with barrel cortex (BC), primary auditory cortex (Aud), and secondary medial visual cortex (V2M) showing the highest and aRSC the lowest increase in the majority of the animals (Supplementary Fig. 4a). There was also a region-dependency in the rate (i.e., slope or derivative) and onset of glutamate concentration change with aRSC showing the steepest slope and the latest elevation onset compared with other imaged regions, especially with BC, Aud, and V2M which showed the lowest slope (Supplementary Fig. 4b) and the earliest onset time in the majority of the animals (Supplementary Fig. 4c). Lastly, the ensemble-wise correlation coefficients averaged across iGluSnFR-Ras animals did not reveal a period of enhanced coordination between hippocampal MUA and aRSC glutamate activity in close vicinity of the ripples (Fig. 2civ-vi; compare with Fig. 2biv).

**Supplementary Fig. 4.**
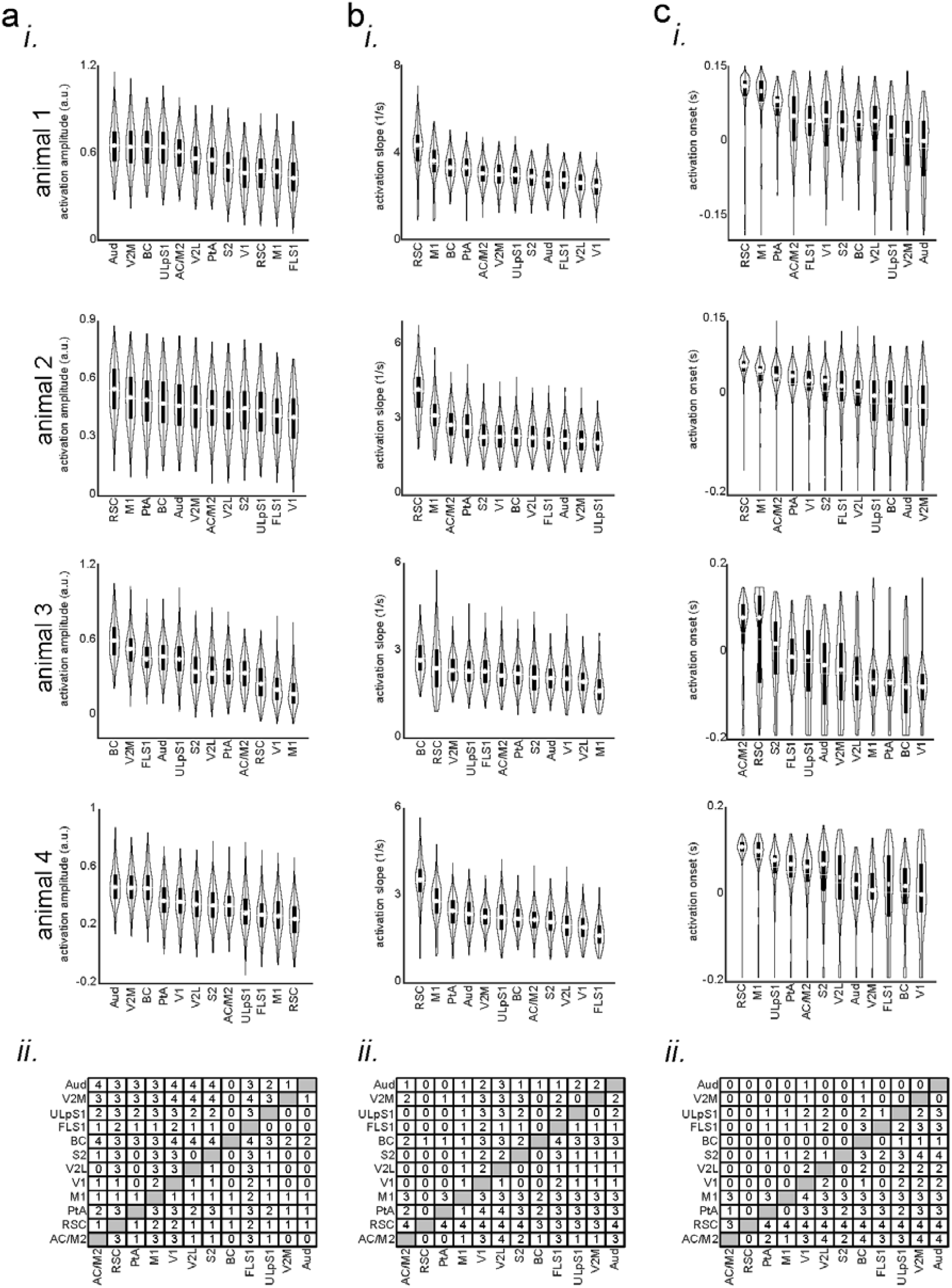
**related to Fig. 2 a (i)** Bootstrap distribution of glutamate activation amplitude (with respect to the pre-ripple baseline) across imaged neocortical regions for individual iGlu-Ras animals.. The regions are sorted according to their distribution mean in descending order (repeated-measure ANOVA with df = 11 and n = 270; from top to bottom: F = 337.9781, df = 11, p-value = 3.6841×10^−262^; F = 304.206, p = 1.3×10^−211^; F = 663.42, p = 0; F = 394.13, p = 4.42×10^−256^). **(ii)** The result of posthoc multiple comparisons following repeated measure ANOVA, pooled across animals (4 animals). The numbers at each entry of the matrix represent the number of mice for which the bootstrap distribution associated with the corresponding row region has statistically significant larger mean than column region. Significance alpha level of 0.05 was used as the threshold for statistical significance. **b-c** The same as (a) but for glutamate activation slope (mean over full-width at half-maximum of derivative of the glutamate signal repeated-measure ANOVA with df = 11 and n = 270; from top to bottom: F = 207.5245, p-value = 3.427×10^−130^; F = 889.69, p = 0; F = 81.93, p = 2.83×10^−76^; F = 249.77, p = 1.35×10^−180^) and glutamate activation onset (timestamp at which the derivative of iGluSnFR signal reaches its half-maximum for the first timep; repeated-measure ANOVA with df = 11 and n = 270; from top to bottom: F = 113.0048, p-value = 7.705×10^−149^; F = 142.93, p = 3.53×10^−187^; F = 86.56, p = 2.53×10^−119^; F = 57.1, p = 5.78×10^−80^). Note that aRSC, compared with all other imaged regions, shows fastest rate of change and latest onset of glutamate activation in at least 3 out of 4 animals.

### aRSC neurons show opposite patterns of peri-ripple modulation

Due to the hyperpolarization of membrane potential in superficial layers of aRSC and the delayed glutamate elevation in the region (Supplementary Fig. 4c: the onset time of RSC activation is larger than zero; Fig. 3a), aRSC neurons may not fire at all during awake ripples. To address this question, we performed two-photon calcium imaging of the neurons in layers II/III of aRSC in Thy1-GCamp mice. Peri-ripple averaging of single cell calcium traces was performed, and the average traces of neurons over the interval -500 ms to +500 ms were clustered into two clusters using the k-means algorithm with correlation coefficient as the similarity metric. Two was the optimum number of clusters according to the silhouette and Calinski-Harabasz criteria. This analysis revealed that there are at least two equally-sized subpopulations of neurons in aRSC; one whose firing is elevated and one whose firing is suppressed during and right after awake ripples (Fig. 3bi). Notably, in the ∼1s-long interval before the ripple centers, the firing of elevated and suppressed sub-populations was suppressed and elevated, respectively (Fig. 3bii). The pre-ripple modulation of the two sub-populations is consistent with the excitatory and inhibitory ramps observed in Chambers et al.^14^. These results show that, despite the presence of significant reduction in population membrane voltage in aRSC, a substantially-large sub-population of neurons increase their firing rate during awake ripples. However, the timing of their firing does not match that of the elevation in glutamate signal (increase in firing precedes glutamate signal elevation). This led us to ask whether the observed glutamate signal consists of components whose timings match those of the observed reduction in voltage signal and the increase firing rate as indicated by calcium imaging. To address this question, we performed singular-value decomposition on the concatenated stack of individual peri-ripple glutamate activity chunks. This method decomposed the stack into components with specific spatial (Fig.3c upper row) and temporal modes. Then, for each component, the corresponding temporal mode was chunked around individual ripples, aligned, and averaged (Fig. 3c lower row). Notably, the first component showed a global post-ripple elevation of glutamate activity whose amplitude was an order of magnitude larger than that in other components. Other components, on the other hand, showed a mixture of elevation and reduction across neocortical regions. These patterns were similar across all the animals. We combined all the components with mixed patterns of modulation (components 2 to 100) and reconstructed the mean peri-ripple glutamate activity across neocortical regions (Fig. 3di-iii). We observed that the mean peri-ripple glutamate activity in aRSC was decomposed into two specific patterns of post-ripple modulation, positive (Figure 3dii; red signal; reconstructed from component 1) and negative (blue signal; reconstructed from components 2 to 100). Interestingly, other regions did not show the negative pattern of modulation (Fig. 3diii). In addition, the timing of the post-ripple negatively-modulated glutamate signal in aRSC matched that of the voltage activity in aRSC (Fig. 3e), which suggests that one of the factors involved in the reduction of voltage in aRSC could be the reduction of endogenous and/or exogenous excitatory glutamatergic input to the region. Also, the onset time of the positively modulated glutamate signal in aRSC was earlier than that of the original signal, which matches better with the timing of aRSC neurons firing presented in Figure 3b. Lastly, the ensemble-wise correlation coefficients averaged across iGluSnFR-Ras animals revealed a period (∼ 0 – 100 ms) of enhanced coordination between hippocampal MUA and negatively-modulated (but not positively-modulated) aRSC glutamate activity (reconstructed from components 2-100) which was absent for sensory regions (Fig. 3div-vii).

**Fig. 3.**
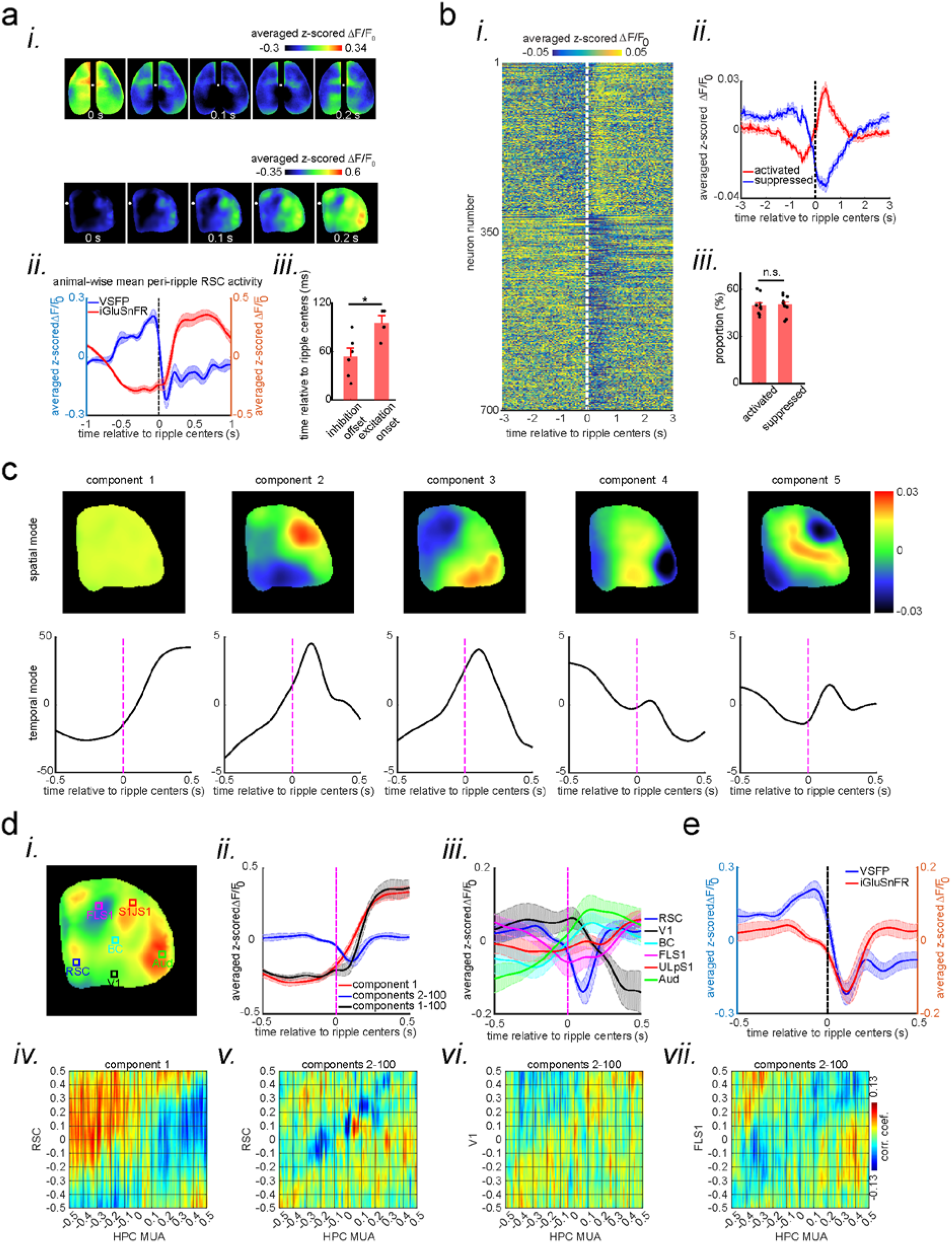
A subpopulation of aRSC neurons fire during awake ripples despite the strong voltage reduction. **a (i)** five frames taken from the montages shown in Fig. 2ai-ii aligned with respect to the ripple center timestamps (zero time). Note the elevation of the glutamate signal as voltage suppression eases. Also note that voltage reduction is strongest and glutamate activation onset is latest in aRSC compared with other regions. **(ii)** time-series representation of the aRSC voltage (blue) and glutamate (red) signals shown in (i). Note that the onset of glutamate activation is around the offset of voltage suppression. **(iii)** Statistical comparison of the voltage suppression offset time in VSFP mice (n=6) and glutamate onset time in iGlu-Ras mice (n=4). There is a statistically significant difference between the two (two-sample t-test; p = 0.02). **b (i)** Average calcium trace (ΔF/F_0_) for individual neurons 3 second before and after ripple centers in a representative Thy1-GCamp animal. The neurons’ calcium traces are grouped into two clusters and are sorted based on their cluster membership. During ripples, the neurons in cluster 1 and 2 show elevation and suppression of calcium signal, respectively. **(ii)** Peri-ripple calcium traces are averaged across neurons in each cluster in each animal and then averaged across 11 animals. The shading represents standard error of the animal-wise mean. **(iii)** Statistical comparison of the proportion of neurons in cluster 1 (activated) and 2 (suppressed). There is no significant difference between the two proportions (paired t-test; p > 0.05). Comparing the results in (aii) and (b-ii) suggests that majority of neurons in clusters 1 and 2 are likely modulated by the excitatory and inhibitory forces applied to aRSC, respectively. **c** Spatial and temporal modes associated with first 5 largest singular values (components) of the concatenated stack of peri-ripple iGluSnFR activity in the representative iGlu-Ras animal presented in Fig. 2aii. Note that the spatial mode of the first component does not show a specific topography and the corresponding temporal mode is dominated with post-ripple elevation of iGluSnFR signal. Also, the amplitude of the first component temporal mode is an order of magnitude larger than that in other components. **d (i)** A representative frame chosen from (a) with 6 ROIs chosen from 6 different neocortical regions. **(ii)** Three animal-wise (n = 4) averages of the reconstructed mean peri-ripple glutamate signals captured from the aRSC ROI in (d-i). The signals were reconstructed using first (red), second-to-hundredth (blue), and first-to-hundredth (black) components in (c). The black signal is the summation of the red and blue ones. Note that the red signal (first component) captured almost all of the elevation seen in the black signal while the blue signal (2-100^th^ components) shows a post-ripple dip. **(iii)** Animal-wise average of the reconstructed mean peri-ripple glutamate signals captured from all the ROIs in (d-i) color-coded according to the ROIs. The signals were reconstructed using second-to-hundredth components. Note that only aRSC shows a post-ripple dip. **(iv)** Ensemble-wise correlation coefficient function of the peri-ripple aRSC glutamate activity (only 1^st^ component) and hippocampal HPC MUA. Rows and columns of the matrices represent time (in seconds) relative to ripple centers. **(v-vii)** The same as (iv) but for glutamate activity of three regions reconstructed from components 2-100^th^. Note the presence and absence of enhanced correlation between aRSC and HPC MUA in the time interval (0,100 ms) in (iv) and (v), respectively. **e** Animal-wise average of mean peri-ripple signals captured from aRSC in all VSFP (blue; n = 6) and iGlu-Ras (red; n = 4) animals. The signals in iGlu-Ras animals were reconstructed from 2-100^th^ components. Note that the timing of the dips in both signals match, suggesting they both represent the same phenomenon.

### The peri-ripple glutamatergic activity is layer-dependent

Given different hypothesized functions for superficial and deep layers in association cortices ^15,16^, we asked whether the patterns of peri-ripple glutamate activity is layer-dependent. To address this question, we conducted wide-field optical imaging with concurrent CA1 LFP/MUA recording in EMX iGluSnFR mice with glutamate indicators expressed in excitatory neurons across all the neocortical layers (as opposed to Ras mice with only superficial layer expression). Qualitatively, the mean peri-ripple glutamate activity and ensemble-wise correlation coefficients in EMX mice (Fig. 4a-b) did not differ from that in Ras mice (Fig. 2aii, c). However, the activity in EMX mice seemed to be shifted to earlier time (compare Fig. 2ciii and 4biii). To probe the potential differences in glutamate activity in Ras and EMX mice, we compared the result of the singular value decomposition analysis. The spatial and temporal modes associated with different components were similar in these two strains. Moreover, the reconstructed signals using 1^st^ and 2-100^th^ components showed the same pattern of positive and negative modulations, respectively (Fig. 4ci-ii). The ensemble-wise correlation coefficients averaged across iGluSnFR-EMX animals revealed a period (∼ 0 – 100 ms) of enhanced coordination between hippocampal MUA and negatively-modulated (but not positively-modulated) aRSC glutamate activity (reconstructed from components 2-100) which was absent for sensory regions (Fig. 4ciii-vi).

**Fig. 4.**
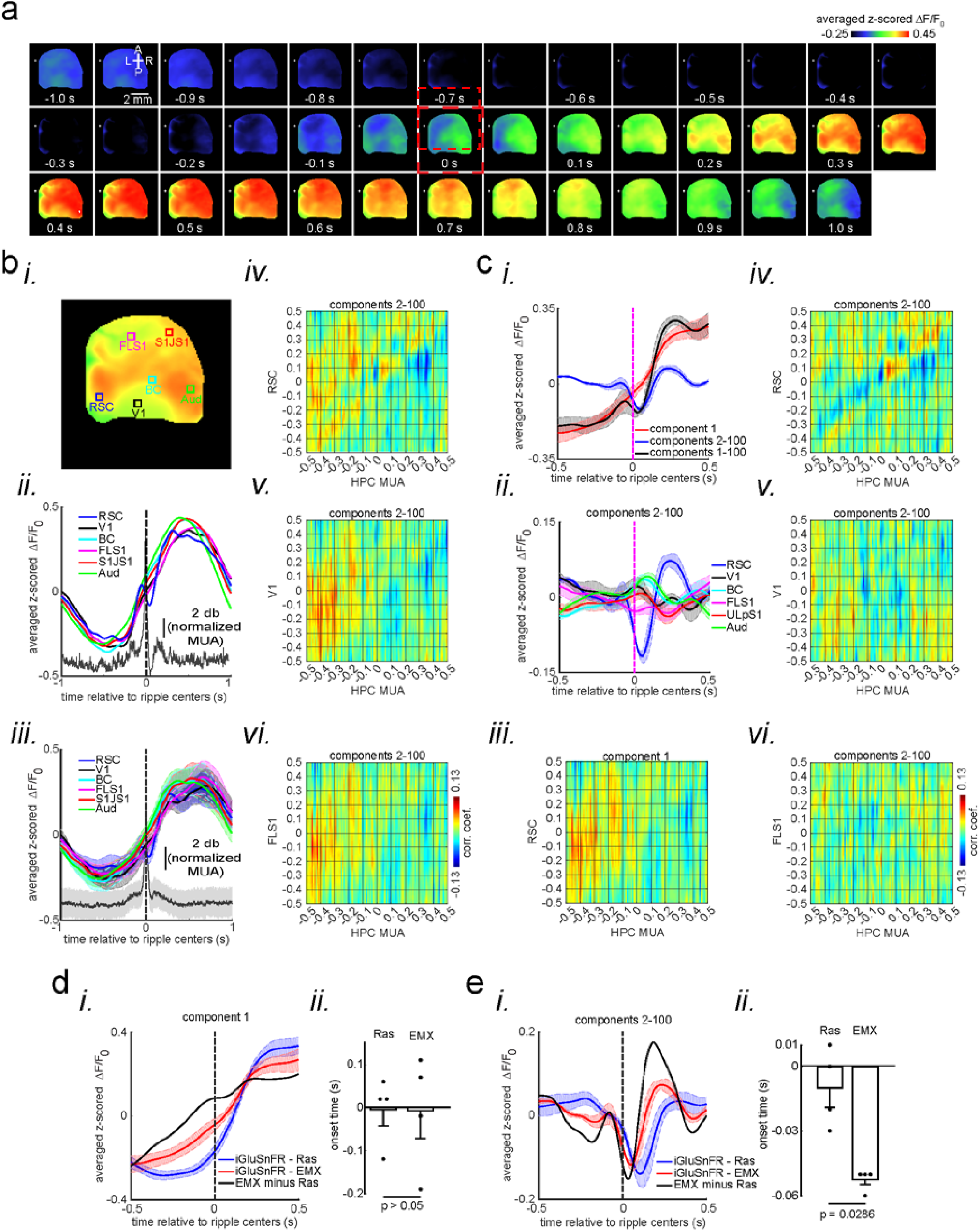
Peri-ripple glutamatergic transmission in neocortical superficial layers is delayed compared with that in deep layers. **a** Montage of average iGluSnFR activity 1 second before and after ripple centers in a representative iGlu-EMX animal. Zero time (red dashed square) represents the timestamp of center of ripples. Note the elevation of glutamate signal across neocortical regions around ripple times. **b (i-ii)** A representative frame chosen from elevation period in (a) along with peri-ripple mean iGluSnFR time-series of six regions of interest chosen from agranular retrosplenial cortex (aRSC), primary visual cortex (V1), and primary forelimb somatosensory cortex (FLS1), primary lip somatosensory cortex (ULpS1), primary barrel cortex (BC), and primary auditory cortex (Aud). The data represented in time-series format is the same data shown in (a). **(iii)** peri-ripple mean iGluSnFR time-series averaged across 4 mice. The shading represents standard error of the mean (SEM). The glutamate signals in iGlu-EMX animals are shifted to left (precede) compared with those in iGlu-Ras animals represented in Fig. 2a-ii and 2c. The black trace represents the mean hippocampal multi-unit activity (HPC MUA). **(iv-vi)** Ensemble-wise correlation coefficient function of the peri-ripple voltage activity of the neocortical regions and hippocampal multi-unit activity (HPC MUA). Rows and columns of the matrices represent time (in seconds) relative to ripple centers. **c (i)** Three animal-wise (n = 4) averages of the reconstructed mean peri-ripple glutamate signals captured from the aRSC ROI in (b-i). The signals were reconstructed using first (red), second-to-hundredth (blue), and first-to-hundredth (black) components. The black signal is the summation of the red and blue ones. Note that the red signal (first component) captured almost all of the elevation seen in the black signal while the blue signal (2-100^th^ components) shows a post-ripple dip. **(ii)** Animal-wise average of the reconstructed mean peri-ripple glutamate signals captured from all the ROIs in (b-i) color-coded according to the ROIs. The signals were reconstructed using second-to-hundredth components. Note that only aRSC shows a post-ripple dip. **(iii)** Ensemble-wise correlation coefficient function of the peri-ripple aRSC glutamate activity (only 1^st^ component) and hippocampal HPC MUA. **(iv-vi)** The same as (b iv-vi) but for signals reconstructed from components 2-100^th^. Note the presence and absence of enhanced correlation between aRSC and HPC MUA in the time interval (0,100 ms) in (iii) and (iv), respectively. **d (i)** Animal-wise (n = 4) average of reconstructed (using 1^st^ component) mean peri-ripple glutamate activity in iGlu-Ras (blue; n = 4) and iGlu-EMX (red; n = 4) animals. **(ii)** The statistical comparison of onset time in iGlu-Ras and iGlu-EMX signals in (i) (two-way ranksum test). **e (i)** Animal-wise average of reconstructed (using 2-100^th^ components) mean peri-ripple glutamate activity in iGlu-Ras (blue) and iGlu-EMX (red) animals. **(ii)** The statistical comparison of onset time in iGlu-Ras and iGlu-EMX signals in (i) (two-sided ranksum test).

Moreover, we did not find a statistically significant difference in amplitude and slope of neither positively-nor negatively modulated signals between Ras and EMX groups. Additionally, although the animal-wise average of positively-modulated aRSC signal showed earlier onset time (Fig. 4di), the statistical comparison between the two groups did not reach the significance threshold of 0.05 (Fig. 4dii). However, the onset time of negatively-modulated signal in aRSC was significantly earlier in EMX than that in Ras group (Fig. 4e). We also estimated the glutamate activity of the deep layers of aRSC by subtracting a scaled version of the Ras from the EMX signal (the black trace in Fig. 4d-ei; 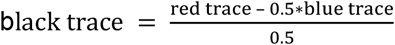). The scaling was performed to accommodate for the potential amount of variance the superficial and deep layers explain in the EMX signal^17^. The estimated positively- and negatively modulated glutamate signals from the deep layers of aRSC showed a shift to an earlier time (to the left). Furthermore, the estimated negatively-modulated signal (black trace in Fig. 4ei) showed a noticeable rebound with a peak around 200 ms after the ripple centers. All in all, these results suggest that deep neocortical layers in aRSC receive glutamatergic modulation earlier than superficial layers do.

## Discussion

In this study, we investigated the peri-ripple activity of the neocortex during the awake state. We utilized voltage, glutamate, and calcium imaging to untangle the activity dynamics in the input, internal, and output levels of neocortical regions, especially in agranular retrosplenial cortex (aRSC). “Input” in this context is considered from the perspective of the dendritic trees occupying the majority of the volume of the superficial layers of the neocortical layers^18,19^. Thus, even self-excitation (i.e., the excitation of the dendrites of a region by the axonal projections originating from the same region) is seen as input in this framework.

The data revealed a reduction in peri-awake-SWR membrane voltage of pyramidal cells with the strongest and fastest hyperpolarization in aRSC. The reduction of membrane voltage could be due to a reduction of the glutamatergic drive or an increase in gabaergic inhibition. We monitored the glutamate delivered to pyramidal cells to test for the first possibility. The mean peri-awake-SWR glutamate concentration did not show a reduction and instead showed a delayed elevation. Moreover, we analyzed the glutamate imaging data further by applying the singular value decomposition (SVD) to see if the glutamate signal was a mixture (multiplex) of negatively- and positively-modulated components. The motivation for investigating the idea of multiplexed glutamate transmission was further fueled by the fact that a significant subpopulation of neurons in aRSC fired despite the significant reduction in voltage as well as seemingly delayed (relative to the SWR centers) glutamate activity in the region. Singular value decomposition (SVD) was able to decompose the potentially-multiplexed glutamate signal in aRSC and to recover the two patterns of modulations. Notably, the timing of these two patterns matched those of voltage reduction and neural firing (measured by calcium activity) in aRSC. It is worthy of note that SVD is agnostic to excitation and inhibition, and it simply captures the maximum amount of variance in the data. Thus, the two patterns of modulations, which resulted from the application of SVD, may not reflect brain mechanisms. However, the coincidence of the timing of these two patterns with those of voltage reduction and neural firing in aRSC suggests that the SVD output, to some extent, reflects truly different neuronal processes.

Even though the peri-awake-SWR glutamate signal was found to be a mixture of rising and falling components, the amplitude (explained variance) of the rising component was larger than that of the falling component. It suggests that the reduction of the glutamate activity in aRSC is not the only factor contributing to the reduction of the voltage signal in the region. Therefore, by exclusion, we suggest that inhibitory input to the superficial layers of aRSC plays a significant role in the reduction of the voltage signal. Moreover, since the majority of the volume of the neocortical superficial layers is filled with dendritic trees^18,19^, dendritic inhibition is probably the major inhibitory process reflected in the reduction of peri-awake-SWR voltage activity.

Another interesting result that came out of the application of SVD on the peri-awake-SWR glutamate activity was the observation that the negatively-modulated component was present only in aRSC and not in other recorded regions while the positively-modulated component was present in all the recorded regions. This observation supports the hypothesis of the presence of peri-awake-SWR inhibition in the neocortex. It is because even though the recorded neocortical regions, except aRSC, did not have a negatively-modulated glutamate activity component, they still showed a reduction in their voltage activity.

Peri-ripple modulation of multiple brain regions, such as ventral tegmental area^20^, subiculum^21,22^, medial prefrontal^8,23,24^, anterior cingulate^25^, and entorhinal cortices^26^, have been observed. Moreover, our data show that multiple neocortical regions, such as the auditory and barrel cortices, express glutamate elevation before aRSC does. Since the majority of these regions project to aRSC, it is plausible that a peri-awake-SWR excitatory input to aRSC comes indirectly from the hippocampus, or even independently from the hippocampus, through these intermediate neocortical regions. In addition, because some aRSC neurons appear to start firing before the timing of SWR centers, it is also plausible that self-excitation may occur in aRSC. In addition, as awake SWRs are involved in planning, and planning could lead to the initiation of movement^27^, another potential source of excitation in aRSC could stem from the subcortical structures involved in motion generation. This could explain as to why EMG tone was detected right after some of the ripples in this study (Supplementary Fig. 1).

According to the current literature, there are at least two potential sources for the peri-ripple inhibitory inputs to aRSC. One option is the long-range inhibitory projections emanating from CA1. Although a majority of tracing studies have been focused on the presence of such projections in the granular RSC (gRSC)^28–30^, it is plausible that they also exist in the agranular RSC (aRSC). The second option is feed-forward inhibition originated from CA1. This mechanism has been reported in gRSC where peri-ripple hippocampal excitatory input activates, via subiculum, the inhibitory interneurons in gRSC which leads to suppression of firing of pyramidal many neurons^31,32^. Similarly, a recent work found that inhibitory interneurons in superficial layers of aRSC increase their firing during awake ripples which, in turn, could suppress the activity of excitatory neurons^14^.

Even though the elevation of neuronal firing is dominant around sleep SWRs, suppression of neuronal firing seems to be an abundant pattern of neuronal modulation around awake ripples. For instance, suppression of neuronal firing around awake hippocampal ripples has been reported in the medial prefrontal cortex^24^ and gRSC^22^. Both of these regions are heavily involved in mnemonic processing, especially through replaying/reactivating the neural traces associated with previous experiences. Given that the fidelity of awake replays is higher than that of sleep ones^12,33^, it could be reasonably speculated that the peri-ripple inhibition plays a role in this higher fidelity. It probably does so via increasing the signal-to-noise ratio by suppressing the interference of the non-mnemonic neuronal populations while the mnemonic representations are being replayed during ripples^24^.

Given the coincidence of peri-ripple inhibition and the negatively-modulated component of the glutamatergic activity in superficial layers of aRSC, it could be deduced that the same coincidence would exist in the deep layers as well. This possibility is supported by the observation that reduction of glutamate signal occurred earlier in deep than superficial layers. In that case, the deep layers would receive a peri-ripple inhibitory force before the superficial layers do. Since the back-projection from the hippocampus mainly target the superficial neocortical layers^16^, the difference in latency of peri-ripple modulation of deep versus superficial neocortical layers could be interpreted from the perspective of the memory indexing theory in the following way: At the time of awake ripples, the hippocampus communicates, via a subspace of neural space, the mnemonic signal as a form of an index code to the superficial layers of RSC. At this time, the deep layers, containing the attributes of past episodic memories, do not receive a mnemonic excitatory drive to avoid interference with the retrieval of index code in the superficial layers^16^. When the index code is retrieved, it is sent to the deep layers for the memory contents to be retrieved, and now the superficial layers are deprived of the mnemonic excitatory drive to avoid any interference with the content retrieval in deep layers. This hypothesized coordinated and sequential retrieval process in superficial and deep layers^16^ might explain the higher fidelity of awake than sleep replays. This is because peri-ripple neocortical inhibition is rare during sleep which implies that the coordination of retrieval of hippocampal content and contextual codes is probably weaker during sleep compared to the awake state.

## Methods

### Animals

6 adult (> 2 months old) male transgenic mice with voltage-sensitive fluorescent protein (VSFP) Butterfly 1.2, expressed in excitatory neurons in neocortical layers II and III, were used for investigating membrane potential dynamics of neocortical regions around awake hippocampal ripples. These mice were generated by crossing the lines Ai78 (Jax023528)^34^, Camk2a-tTA (Jax007004), and Rasgrf2-2A-dCre (Jax022864).

4 adult (> 2 months old) female transgenic mice with fluorescent glutamate indicator (iGluSnFR^35^)^36^, expressed in excitatory neurons in neocortical layers II and III, were used for investigating dynamics of excitatory synaptic input to the neocortical regions around awake hippocampal ripples. These mice were generated by crossing the lines Ai85 (Jax026260), Camk2a-tTA (Jax007004), and Rasgrf2-2A-dCre (Jax022864). These mice are called iGlu-Ras in this work.

4 adult (> 2 months old) male transgenic mice with fluorescent glutamate indicator (iGluSnFR), expressed in excitatory neurons in all neocortical layers, were used for investigating dynamics of excitatory synaptic input to the neocortical regions around awake hippocampal ripples. These mice were generated by crossing the lines Ai85 (Jax026260), Camk2a-tTA (Jax007004), and Emx1-Cre (Jax005628). These mice are called iGlu-EMX in this work.

11 Thy1-GCaMP6s female mice with fluorescent calcium indicator, expressed in excitatory neurons across all neocortical layers, were used for investigating spiking dynamics of neurons in layers II/III of agranular retrosplenial cortex around awake hippocampal ripples.

Mice were housed in groups of two to five under a 12 hr light-dark cycle. Mice were given ad libitum access to water and standard laboratory mouse diet at all times. After head-plate/electrode implantation surgery, the mice were single-housed. The animal protocols were approved by the University of Lethbridge Animal Care Committee and were in accordance with guidelines set forth by the Canadian Council for Animal Care.

### Surgeries for wide-filed voltage and glutamate imaging experiments

On the days of surgery on mice, used in wide-field imaging experiments, subcutaneous injection of buprenorphine (0.5 gr/Kg) was delivered half an hour before the surgery started. Animals were then anesthetized with isoflurane (1–2% mixed in O_2_). After reaching the desired depth of anesthesia, the following steps were performed: (1) the skull skin was removed. (2) Hippocampal LFP electrode was implanted. (3) A head-plate was implanted. (4) The muscles covering the lateral portion of the skull (on top of secondary somatosensory and auditory cortices) were removed. This step was performed only for the glutamate imaging experiments where a unilateral imaging window was used. This step allowed us to image activity of secondary somatosensory and auditory cortices. (5) The skull was covered with a thin and transparent layer of the metabond (Parkell, Inc). (6) The skull was covered with a glass coverslip. An additional bipolar electrode was implanted in the neck muscles for recording EMG activity. Animals were allowed to recover for two weeks before recordings started.

### Surgeries for two-photon calcium imaging experiments

On the days of surgery on mice, used in two-photon calcium imaging experiments, subcutaneous injection of buprenorphine (0.5 gr/Kg) was delivered half an hour before the surgery started. Animals were then anesthetized with isoflurane (1–2% mixed in O_2_). After reaching the desired depth of anesthesia, the following steps were performed: (1) the skull skin was removed. (2) A small craniotomy was performed to remove portion of the skull covering agranular retrosplenial cortex. (3) The exposed part of the brain was covered with a glass coverslip. (4) Hippocampal LFP electrode was implanted. (5) A head-plate was implanted. (6) An additional bipolar electrode was implanted in the neck muscles for recording EMG activity. Animals were allowed to recover for two weeks before recordings started.

### Hippocampal LFP recording

Teflon coated 50 µm stainless steel wires (A-M Systems) were used to make bipolar hippocampal LFP electrodes. The two tips of the electrode were separate around 0.5 mm so that the tips could record from two different depths. To implant the electrode for wide-field imaging experiments, a hole was drilled on the right hemisphere skull about 2.6 mm lateral to the midline and tangent to the posterior side of the occipital suture. Then, the electrode was gradually lowered through the hole at an angle of 57 degrees with respect to the vertical axis (the axis perpendicular to the surface on which the stereotaxic apparatus was sitting). The electrode signal was being continuously monitored both visually and audibly. Lowering the electrode was stopped as soon as a dramatic increase in the spiking activity was heard and observed for the second time near the calculated coordinate (angle = 57 degrees, depth = ∼1.75 mm) for the pyramidal layer of the dorsal CA1. In this way, we ensured that the upper and lower tips of the electrode were placed in and beneath the pyramidal layer of the dorsal CA1, respectively. The electrode was fixated on the skull using Krazy Glue and dental cement. For the two-photon imaging experiments, a similar electrode implantation procedure was used. The only difference was that the electrode was lowered perpendicular to the surface of the brain until it reached the pyramidal layer of the dorsal CA1. In all the experiments, the hippocampal electrodes were implanted in the right hemispheres ipsilateral to the imaging window. The electrode signals were amplified using a Grass A.C. pre-amplifier Model P511 (Artisan Technology Group, IL) and digitized using a Digidata 1440 (Molecular Device Inc, CA) or National Instruments data acquisition system.

### Glutamate imaging

Blue-light-emitting diodes (Luxeon K2, 473 nm, Quadica Developments Inc, Lethbridge, Alberta) augmented with band-pass filters (Chroma Technology Corp, 467–499 nm) were used to excite iGluSnFR indicators. The fluorescence emission from iGluSnFR was filtered with a 520–580 nm band-pass filter (Semrock, New York, NY) and collected as 12-bit images at 100 Hz using a CCD camera (1M60 Pantera, Dalsa, Waterloo, ON) and an EPIX E4DB frame grabber controlled with XCAP 3.7 imaging software (EPIX, Inc, Buffalo Grove, IL). To reduce the effect of large neocortical blood vessels in imaging quality, the lens was focused into the neocortex to a depth of ∼1 mm. We also recorded the iGluSnFR signal in response to different periphery stimulation under urethane anesthesia^13,37^ to functionally map the center of the hind-limb somatosensory, fore-limb somatosensory, auditory, visual, and barrel cortices.

### Voltage imaging

Blue-light-emitting diodes (Luxeon K2, 473 nm, Quadica Developments Inc, Lethbridge, Alberta) augmented with band-pass filters (Chroma Technology Corp, 467–499 nm) were used to excite the Butterfly indicator. FF580-FDi01-25×36 dichroic mirror was used for mCitrine/mKate2 emission light separation before getting filtered using a 528–555 nm and 582-602 band-pass filters (Semrock, New York, NY), respectively. The filtered signals were collected as 12-bit images at 100 Hz using two CCD cameras (1M60 Pantera, Dalsa, Waterloo, ON) and EPIX E8 frame grabber controlled with XCAP 3.7 imaging software (EPIX, Inc, Buffalo Grove, IL). To reduce the effect of large neocortical blood vessels in imaging quality, the lens was focused into the neocortex to a depth of ∼0.5 mm. We also recorded the Butterfly FRET signals in response to different periphery stimulation under urethane anesthesia to functionally map the center of the hind-limb somatosensory, fore-limb somatosensory, auditory, visual, and barrel cortices.

### Two-photon calcium imaging

Two-photon calcium imaging was conducted via a Bergamo II multi-photon microscope (THORLABS). Ti:Saphire pulsed laser (Coherent) with wavelength of 920 nm and power of ∼80 mW (measured at the tissue) was used to excite the calcium indicators. Scaning of the field of view was done by Galvo-Resonant X-Y mirrors. A 16x water-immersion objective lens (Nikon) with numerical aperture of 0.8 was used for imaging. The emitted light from calcium indicators was collected via a GaAsP photomultiplier tube (Hamamatsu). The field of view size was 835 × 835 µm and frames were captured at spatial resolution of 800 × 800 pixels and temporal resolution of 19.6 Hz. The depths of imaging was aimed between 110 and 190 µm (layers II/III).

### Preprocessing of Butterfly (VSFP) imaging data

We followed the ratiometric procedure, described in Carandini et al.^38^, with a modification to preprocess the VSFP data and obtain an estimate of the membrane potential at each pixel. The modification was that we used time-varying quantities for 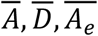, and 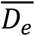 by calculating trends of these signals using the local regression method. We made this adjustment because we were working with spontaneous neocortical activity recorded over a long period of time while, in the Carandini et al.^38^, peri-stimulus activity over a short interval of time (couple of seconds) was analyzed.

### Preprocessing of iGluSnFR imaging data

First, low-rank reconstruction of the stack of frames, obtained via iGluSnFR imaging, was performed by applying singular-value decomposition and taking the components with the greatest associated singular values^39^. Next, for each pixel in the imaging window, a time-varying baseline (F_0_) for the iGluSnFR signal (F) was calculated. Baseline calculation was performed by applying the *locdetrend* function in the Choronux toolbox^40^ (http://chronux.org/) to fit a piecewise linear curve to the pixels’ time series using the local regression method. The calculated baseline signal (F_0_) was then subtracted from the raw signal (F), and the difference signal was divided by the baseline values at each time point (Δ*F*/*F*_0_). At the end, a band pass (0.5–6 Hz) FIR filter was applied on the Δ*F*/*F*_0_ signal for each pixel.

### Preprocessing of two-photon calcium imaging data

The preprocessing of two-photon calcium imaging data was conducted via Suite2p pipeline implemented in Python 3^41^. The signals from the detected candidate neuronal ROIs as well as the geometric shape the ROIs were visually inspected to screen for non-somatic compartments. The neuropil component of the calcium traces, estimated by Suite2p, was multiplied by 0.7 and subtracted from the traces^41^.

### SWRs detection

The raw hippocampal LFP was down-sampled to 1 kHz, filtered between 110 to 250 Hz (ripple-band) using real-valued Morlet wavelet implemented in MATLAB (MathWorks). The ripple power signal was generated by rectifying and smoothing the ripple-band filtered signal. Smoothing was performed using a rectangular window with a length of 8 ms. SWRs were identified when the ripple power signal passed the detection threshold defined by the mean plus a multiple of its standard deviation. The numerical value of the standard deviation multiplier was adjusted manually for each animal. A lower threshold (75% of the detection threshold) was used to estimate the onset and offset of each SWR. Detected events were further screened by applying a duration threshold. The timestamp of the largest trough between the onset and offset times of each detected event is referred to as the event center. At the end, events with centers less than 50 ms apart were concatenated.

### Exclusion of ripples based on EMG activity

To ensure that the peri-ripple neocortical activity was least affected by movement-related brain activity, the ripples with above-threshold EMG activity within ±500 ms were excluded from all the analyses used in this study. The exclusion threshold was manually chosen for each animal.

### Multi-unit activity (MUA) calculation

MUA signal was calculated from hippocampal LFP using a similar method reported before^42^. Briefly, the hippocampal LFP signal were filtered above 300 Hz, rectified and smoothed with a rectangular window with the length of ∼3 millisecond. The resultant signal was called MUA in this work.

### Z-scoring peri-ripple neocortical activity

The z-scoring of peri-ripple traces/frames was performed against a null distribution. The null distribution was obtained from traces centered round random timestamps which did not necessarily correspond to those of ripple centers, and the random timestamps were generated by randomly permuting the intervals between the successive ripple centers. All the individual peri-ripple traces/frames were z-scored against the null distribution before being analyzed further.

### Calculating the number of bootstrap draws and sampling size

The number of bootstrap draws for voltage (i.e., 193) and for glutamate (i.e., 270) imaging data was calculated to achieve the statistical power of 0.8 at significance level of 0.05 and effect size of 0.25 for a repeated-measure ANOVA design with 8 and 12 groups (i.e., regions), respectively. The power analysis was performed using the G*Power software^43^. 50 was chosen as the sampling size at each bootstrap draw. This number was chosen since the correlation coefficient between the average of the bootstrap draws and the average of the whole peri-ripple ripples plateaued around this number in all the animals.

### Calculating amplitude, slope, and onset of activation and/or deactivation

For voltage peri-ripple mean traces, to report all the quantities as a positive number, the traces were inverted by multiplying a negative one to them. Deactivation amplitude was calculated as the difference between maximum value of the trace in the interval [0,200 ms] and the baseline value. The baseline value was calculated as the mean of the trace in the interval [-200 ms,0]. To calculate the onset and offset of the deactivation, the maximum value of the derivative (rate of change) of the voltage traces in the interval [-200 ms,200 ms] was calculated. Onset and offset of deactivation were defined as the timestamps at which the derivative signal reaches half of its maximum value before and after the maximum value timestamp, respectively. The slope of deactivation was defined as the average slope of the voltage traces between their onset and offset times. Finally, the pre-ripple amplitude of the voltage traces were calculated by averaging the non-inverted trace values between the timestamps of the half-maximum value. For glutamate peri-ripple mean traces, all the calculations are the same as those applied to the voltage imaging data except the signal inversion at the very first step was not performed.

### Calculating ensemble-wise correlation coefficient function

Peri-ripple activity of each region and peri-ripple MUA could be conceived of as random processes for which we have observed multiple realizations. The observed realizations of these two random processes could be arranged in the matrix form as A(*r,t*) and B(*r,t*), respectively, where *r* and *t* represent *r*-th ripple and *t*-th time point with respect to the *r*-th ripple center. Note that all the observed realizations are aligned with respect to the ripple centers. The cross-correlation coefficient function, C(*t*_*1*_,*t*_*2*_), between random processes A and B could be estimated by calculating the sample correlation coefficient between A(:,*t*_*1*_) and B(:,*t*_*2*_). In other words, C(*t*_*1*_,*t*_*2*_) equals the correlation coefficient between *t*_*1*_-th column of A and *t*_*2*_*-th* column of B.

### Clustering the aRSC calcium traces

The z-scored (against the null distribution) peri-ripple calcium traces in the interval [-500ms, 500ms] were further z-scored with respect to their mean and standard deviation and then fed into the k-means algorithm implemented as the built-in function *kmeans* in MATLAB (MathWorks). Correlation was chosen as the distance metrics.

### Statistical tests

All statistical tests in this study were performed using MATLAB built-in functions. Repeated-measure ANOVA with Greenhouse-Geisser correction for sphericity was performed for testing the hypothesis that there was a region-effect in any of the features (i.e., amplitude, slope, onset/offset time) of the peri-ripple traces. This analysis was followed by performing multiple comparisons with Tukey-Kramer correction. For all the two-group comparisons, two-sample two-sided t-test was used.

## Author contributions

Conceptualization, J.KA., B.L.M., M.H.M.; Methodology, J.KA., Z.J., M.H.M..; Investigation, J.KA, Z.R.; Formal Analysis, J.KA.; Experimental models, T.K.; Writing – Original Draft, J.KA; Writing – Review & Editing, J.KA., Z.R., T.K., B.L.M., and M.H.M.; Funding Acquisition, M.H.M., B.L.M.; Resources, B.L.M and M.H.M.; Supervision, M.H.M.

## Acknowledgment

This work was supported by Natural Sciences and Engineering Research Council of Canada (grant no 40352 & 1631465 to MHM and BLM respectively), Alberta Innovates (MHM and BLM), Alberta Prion Research Institute (grant no. 43568 to MHM), and Canadian Institute for Health Research (grant no 390930 & 156040 to MHM and BLM MHM respectively), National Science Foundation (MHM, BLM), and USA Defense Advanced Research Projects Agency (grant no HR0011-18-2-0021 to BLM). The authors also thank Di Shao and Behroo Mirza Agha for animal breeding, and J. Sun for surgical assistance. The authors thank Hongkoi Zeng from the Allen Institute for Brain Science for providing the Emx-cre, Camk2a-tTa, and Ai85 mice as a gift.

